# HSF1 Activator Azadiradione Ameliorates Parkinson’s Disease and Extends Lifespan in Preclinical Models: Analysis of Underlying Molecular Mechanism

**DOI:** 10.1101/2025.04.09.647992

**Authors:** Hossainoor Rahaman Sareng, Naibedya Dutta, Suvranil Ghosh, Upama Chowdhury, Vinod K. Nelson, Mohit Prasad, Subhash C. Mandal, Atin K. Mandal, Mahadeb Pal

**Author notes:** Equal Contribution (Hossainoor Rahaman Sareng, Naibedya Dutta).

## Abstract

Parkinson’s disease (PD) affects millions worldwide, with no efficient therapy currently available. A major cause of the initiation and progression of this degenerative disease is the dysfunctional cellular protein quality control system (PQC), leading to the accumulation of toxic protein aggregates in neurons. We previously reported azadiradione (AZD), a small molecule (MW 451 Da), as a potent inducer of heat shock factor 1 (HSF1) activity, which could alleviate cellular toxicity induced by misfolded proteins by upregulating the levels of inducible molecular chaperones and proteasome activity. Here, we show the multifaceted effect of AZD in enhancing the capacity of PQC machinery in cells, fruit flies, and a PD mouse model. AZD activated HSF1 by promoting its phosphorylation at S326 through MEK. In parallel, AZD boosted protein degradation through increased chymotrypsin-like proteasome activity, upregulation of the ubiquitin ligase CHIP. AZD induced autophagy, marked by elevated levels of Beclin 1, ATG7, and ULK1 phosphorylation at S555, along with mTORC1 inhibition via AMPK activation. Surprisingly, the calorie restriction pathway was also upregulated upon AZD treatment, as demonstrated by the enhanced phosphorylation of FOXO3 and FOXO1, along with increased activity of their target enzymes SOD and catalase. Notably, AKT activity was also suppressed in AZD-treated cells. In vivo, AZD improved motor function, dopaminergic neuron survival, and tyrosine hydroxylase activity in an MPTP-induced mouse model of PD, and extended lifespan in Drosophila without compromising fertility or mobility. These findings highlight AZD as a promising therapeutic candidate for restoring PQC and mitigating PD pathology.

## Introduction

Neurodegenerative disorders (NDs), such as Parkinson’s disease (PD), Alzheimer’s disease, and amyotrophic lateral sclerosis (ALS), pose significant challenges to human health due to the limited therapeutic options currently available. These disorders primarily affect individuals aged 60 and older, leading to progressive declines in cognitive and motor functions, and often advancing to dementia [1-5]. According to the World Health Organization (WHO), the global population aged 60 years and older is expected to exceed 2 billion by 2050—more than double the current number [6-8] . The Global Burden of Disease study projects that NDs will become one of the leading causes of death worldwide by 2050 [3].

Among the NDs, the incidence of PD is increasing most rapidly [9, 10]. According to the Parkinson’s Foundation, approximately 90,000 Americans are diagnosed with PD each year, representing a 50% increase compared to previous estimates. However, data on the incidence and prevalence of PD and other NDs in India remain limited due to challenges in healthcare infrastructure, access to medical care, and reporting systems. The incidence of PD varies significantly across different communities in India, with an average rate of approximately 70 per 100,000 people. Notably, the Parsi community in Mumbai experiences a much higher incidence rate of PD—about 328 per 100,000 people—making it one of the highest rates worldwide [11].

PD manifests through a combination of motor and non-motor symptoms. Motor symptoms include tremors, bradykinesia (slowed movement), muscle rigidity, postural instability, and impaired balance and coordination. Non-motor symptoms include cognitive decline, dementia, constipation, and sleep disturbances [12]. The key pathological features of PD are the loss of dopaminergic neurons in the substantia nigra pars compacta of the midbrain and the accumulation of protein aggregates, such as Lewy bodies and Lewy neurites. These aggregates primarily contain alpha-synuclein, Parkin, and ubiquitinated proteins [13]. A critical aspect of PD pathogenesis is the dysfunction of protein quality control (PQC) mechanisms, including the heat shock response (HSR), ubiquitin-proteasome system (UPS), and autophagy-lysosomal pathway (ALP). Impairments in these processes are believed to contribute significantly to the development of PD and other neurodegenerative disorders [14-16].

Heat Shock Factor 1 (HSF1), a transcription activator of ∼54 kDa, is the master regulator of the heat shock response (HSR). Under normal conditions, HSF1 exists in the cytoplasm as an inactive monomer in association with a repressive complex consisting of chaperones such as HSP70, HSP90, and TRiC/CCT [17-19]. However, upon exposure to cellular stress, HSF1 dissociates from the repressive complex, forms a homotrimer, and translocates to the nucleus, where it binds to its recognition sequence motif called the heat shock element (HSE) in the regulatory regions of target genes. HSE is composed of repeating units of the nucleotide sequence 5’-nGAAn-3’ [20]. These genes encode inducible HSPs and various proteases, which help refold misfolded proteins into their native, functional conformations or degrade them into constituent amino acids via the ubiquitin-proteasome system (UPS) or the autophagy-lysosomal pathway (ALP), respectively [21-23] .

Post-translational modifications, such as phosphorylation by various protein kinases—including MEK, ERK, mTOR, AMPK, and ULK1—along with acetylation/deacetylation mediated by CBP and SIRT1, play crucial roles in regulating HSF1 activity [24] . MEK promotes phosphorylation of HSF1 at serine 230 [25] and serine 326 during activation [26-28].

In addition to regulating inducible protein chaperones such as HSP70, HSF1 also controls the expression of certain regulators in the UPS and ALP pathway [24, 29]. HSP70, besides chaperoning its client proteins, facilitates the proteasomal degradation of misfolded substrates through the carboxy terminus of HSP70-interacting protein (CHIP), an E3-ubiquitin ligase [30, 31]. The UPS and ALP are responsible for the elimination of distinct types of misfolded proteins, protein aggregates, and dysfunctional organelles generated during cellular stress [32]. The proteasome (UPS) primarily degrades soluble and short-lived regulatory proteins, while autophagy targets larger protein aggregates and damaged organelles, ensuring proper turnover of cellular components [29, 33]. Disruption of either pathway can lead to protein aggregation and contribute to disease pathology, particularly in neurodegenerative disorders [34, 35].

Earlier, we isolated AZD from the seeds of *Azadirachta indica*, a plant well known as ‘neem’ for its many medicinal properties. We demonstrated that AZD ameliorates toxicity caused by polyglutamine (polyQ)-rich protein aggregation in cells as well as in fruit flies. AZD was uniquely shown to enhance the binding of HSF1 to its recognition sequence, HSE, by direct physical interaction in vitro and in cells, correlating with the increased expression of its target genes, such as the inducible HSPs [36]. Notably, AZD does not induce oxidative imbalance in the cell, unlike many other plant-based small molecules reported previously [37]. AZD was efficient in ameliorating Huntington’s disease in mice [38]. To date, AZD is the only HSF1 activator that appears to function in this way. However, the detailed understanding of AZD-mediated HSF1 activation and its potential to ameliorate PD in cellular and preclinical models remains to be tested.

Given the demonstrated role of HSPs/HSR in cellular aging, we tested how AZD, an apparent small molecule mimic of HSF1, influences lifespan in an experimental animal. *Drosophila melanogaster* (fruit fly) was used as a model in this test because of its relatively short lifespan [39].

## Materials and Methods

### Cell culture

Mouse neuroblastoma 2A (Neuro-2A) and human embryonic kidney 293 (HEK293) cells obtained from ATCC were cultured in DMEM (Gibco) supplemented with 10% fetal bovine serum (Gibco North American, heat inactivated), L-glutamine (1 mM), penicillin (50 μg/ml), streptomycin (50 μg/ml), amphotericin B (2.5 μg/ml), gentamycin (50 μg/ml) and non-essential amino acids, and incubated at 37ºC in a humidified incubator under 5% CO2. Cells were split at around 90% confluency to the fresh media, and treated at 70% confluency [40].

### Short hairpin RNA construct preparation and transfection assay

Short hairpin construct of FOXO3 and AMPK was cloned in PLKO.1 puro vector. siRNA construct of HSF1 and CHIP were obtained from Genex, India. Transfection experiments were carried out at 70% confluency using lipofectamine 2000 as described [41]

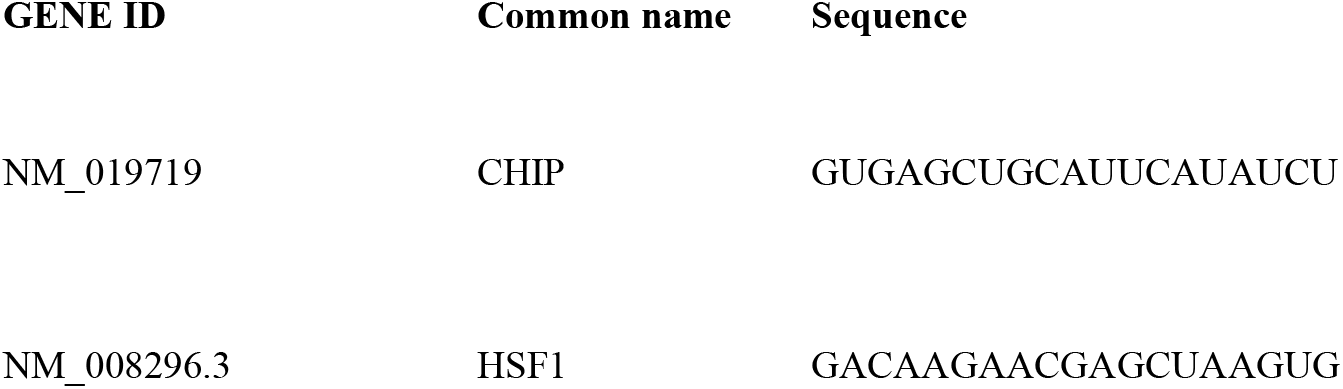

### Preparation of whole-cell lysate

Cells grown in plates/flasks, after treatment were harvested in ice cold PBS through scrapping. The whole cell lysates (WCL) was prepared in a lysis Buffer [20 mM Tris-HCl pH 7.5, 1% Triton X-100, 300 mM NaCl, 5% glycerol, 1 mM phenylmethyl sulfonylfluoride (PMSF) supplemented with 10 μg/ ml leupeptin, 10 μg/ml aprotinin, 20 mM sodium fluoride and 20 mM sodium orthovanadate]. The protein concentration in a WCL was measured using Bradford assay reagent (BioRad) with BSA as standard [40]

### Western blot assay

Proteins in the WCL (30 µg/sample) resolved through SDS-PAGE (acrylamide: bisacrylamide::29:1) were transferred to PVDF membrane pre-fixed in methanol through wet transfer method. Then the membrane was incubated for 1 h at RT in blocking solution [5% BSA dissolved in PBST (PBST:137 mM NaCl, 2.7 mM KCL, 10 mM Na2HPO4, 1.8 mM KH2PO4 pH 7.4, supplemented with 0.1% Tween 20) before incubating with an appropriate dilution of desired primary antibody for overnight at 4ºC. The membrane was washed with PBST, incubated with HRP conjugated secondary antibody diluted (1:3000) in PBST for 1 h at RT. The membrane was washed, and finally signal was developed by enhanced chemiluminescence method by ECL Clarity Bio-Rad and detected with a chemidoc (BioRad) [36]

### Proteasome activity assay

Proteasome activity in the WCL was measured using Proteasome 20S Activity Assay Kit (SIGMA-MAK172). This kit uses LLVY-R110 substrate as a fluorogenic indicator for protease activity. Proteases cleave in LLVY-R110 to release fluorogenic R110 (excitation at 490 nm, emission at 525 nm). Bortezomib (100 nM) a proteasome inhibitor was used as a positive control [41]

### Antioxidant enzyme activity assay

WCL were prepared from mouse neuroblastoma 2A (Neuro-2A) cells pre-treated as appropriate for 24 h. Superoxide dismutase (SOD) activity level was estimated in the WCL by its formazan formation measured at 560 nm [42]. Catalase (CAT) activity was estimated by measuring the extent of breakdown of H_2_O_2_ at 240 nm [43]

### Immunofluorescence assay

Cells were grown on round grease-free cover slips to 50% confluency. After desired treatment, the cells were washed with PBS before fixing in 4% paraformaldehyde for 20 minutes at 37ºC. Cells were washed with PBS, incubated with 1% triton X-100 for 15 mins at RT followed by several washes with PBS. After blocking with 2% BSA made in PBST for 1 h at RT, cells were incubated with primary antibody at 1:100 diluted in 1% BSA-PBST for overnight at 4ºC in a humidified chamber. Next day cells were washed with PBS before incubating with fluorescence conjugated secondary antibody for 2 h in the dark. The cells on the coverslips were then washed with PBS before mounted on a slide with VECTASHIELD mounting medium for capturing the images under confocal microscope (Leica) [41]

### Animal care and MPTP treatment

Male Swiss albino mice (from 6 to 8-week-old) were obtained from Bose Institute animal facility and housed for a week in a controlled temperature (23+/-1°C), 12-h light/dark cycle and with ad libitum access to food and water for acclimatisation. Then the mice were randomly sorted into four groups (A to D), with each group having 5 mice (n=5). To develop Parkinsonism, MPTP was injected (in group C, D) subcutaneously 3 times in a week for three weeks at a dose of 10 mg/kg body weight. The control group (group A) received a similar treatment with vehicle control (DMSO). After completion of MPTP administration, group B and group D were administered with AZD through IP injection (20 mg/kg body weight for 20 days with every alternate days)[41]. Mice experiments were carried out with prior permission from Bose Institute animal ethics committee (IAEC/BI/88/2018).

### Immunohistochemistry of mice brain tissue samples

Transcardial perfusion was done in anesthetized mice with PBS and 4% paraformaldehyde (1:1). Brains after isolation were immediately fixed in 4% paraformaldehyde diluted in PBS. Brains embedded in paraffin block were sliced into 30 nm thin section on a freezing microtome and affixed onto lysine coated slides. The paraffin embedded sections were deparaffinised by washing with xylene (2 times, 3 minutes each). The slides were then washed sequentially with xylene:ethanol (1:1), 100% ethanol, 90% ethanol, 70% ethanol, and 50% ethanol (each step for 3 minutes). Finally, the slides were rinsed with cold running tap water. Immunohistochemistry with anti-tyrosine hydroxylase (TH) and HSF1 antibodies were performed using IHC staining kit procured from Vector Laboratories [41]

### Behavioural tests

Body weight of mice was recorded alternative day from the start of the treatment. To test the motor activity, grip test and footprint studies were carried out. For footprint test, animals were trained in a dark tunnel (10*10*40) for 2-3 days before test. Then at the time of test forelimbs were dipped in red ink and hind limbs were dipped with blue inks. Foot marks of mice were recorded on a white paper placed on the floor of the tunnel. By measuring the distance between each step on the same side of the body, stride lengths were determined. We also recorded the total no of footprints inside the tunnel (first two prints are excluded and only steps perform with a straight line with a regular velocity were recorded). For grip test, we recorded the duration the mice were able to hang holding the cage ring with their forelimbs [41] .

### Lifespan assay

A stock of Canton S wild type flies (*Drosophila melanogaster*) were maintained at 25°C with 12 h light/dark cycle on cornmeal diet as described[44]. We choose female fly for the study because of their large body size and reproductive fitness. Two hundred age-matched flies in total were taken for the study. Five vials each 20 flies were kept in media supplemented with AZD while other 5 vials were kept in media supplemented with the vehicle (DMSO/control). Flies were transferred into a fresh vial carrying appropriate growth media every alternative day with the number of dead flies recoded. Transfer to vials continued till the last fly died in the control vials. (Eclosion rate) 30 embryos/larvae were transferred in each vial (in triplicate) carrying the growth media supplemented with different concentration of AZD or vehicle to record the number of flies recorded after 10 to 15 days eclosed in the vials [45]

### Reproductive fitness assay

Age-matched single virgin male and female flies were kept together in the vials in triplicate. The no of embryo/larvae in an embryo collection cage were counted for 24 h under the light microscope [46]

### Negative geotaxis assay

Age-matched female flies (20 flies/vial) of both control and AZD-fed were taken. Flies were transferred into clean transparent graduated measuring cylinder (length 30 cm, diameter 3 cm). After 10 minutes of acclimatization, the cylinder was gently tapped to collect the flies to the bottom. After 10 seconds, the no of flies climbed above 15 cm were counted both in the control and experimental cylinders. The process was repeated three times for each set [47] .

### Transcriptome analysis

Mouse neuroblastoma cells (N2A), grown to 70% confluency in 35 mm plates, added with the indicated concentration of AZD were incubated for 24 h or HS for 30 mins at 39°C. The cells were collected in ice cold 1X PBS in 1.5 ml centrifuge tubes. After a PBS wash the RNA was isolated by TRIzol method (Gibco). Following one round of ethanol precipitation RNA concentrations in the samples were quantified using nanodrop (Thermo). Three biological replicates were measured per condition. Library preparation, and RNA sequencing and analysis were performed at Eurofin Genomics India Pvt. Ltd. Differential gene expression analysis was performed using Cuffdiff version 2.2.1. The analysis was carried out comparing the commonly expressed genes including the genes of our interests reported between the control and treated samples. FPKM values were used to calculate the log fold change as log2 (FPKM_Experimental/FPKM_Control). Log2 Fold Change (FC) values greater than zero were considered up-regulated whereas less than zero were down-regulated along with P-value threshold of 0.05 for statistically significant results.

## Results

### Azadiradione induces activatory phosphorylation of HSF1 through MEK

Different post-translational modifications at various residues are thought to help functional activation of HSF1 [24]. Among these modifications, phosphorylation at S326 is considered as a hallmark of the functionally active state of HSF1 [48]. This phosphorylation can be controlled either by mTOR or MEK depending on the cellular context [Fig. 1*A*][28, 48]. Our lab has previously reported that azadiradione (AZD) could induce cellular HSF1 function including its DNA binding capability [36]. Our test revealed that AZD upregulated phosphorylation of HSF1 at S326 in HEK293 cells [Fig. 1*B*]. Interestingly, in AZD treated cells, increased phosphorylation of MEK was observed suggesting its activation, while reduced phosphorylation of S6K, a downstream target of mTORC1proposing mTOR inhibition [Fig.1*B*].

**Figure 1.**
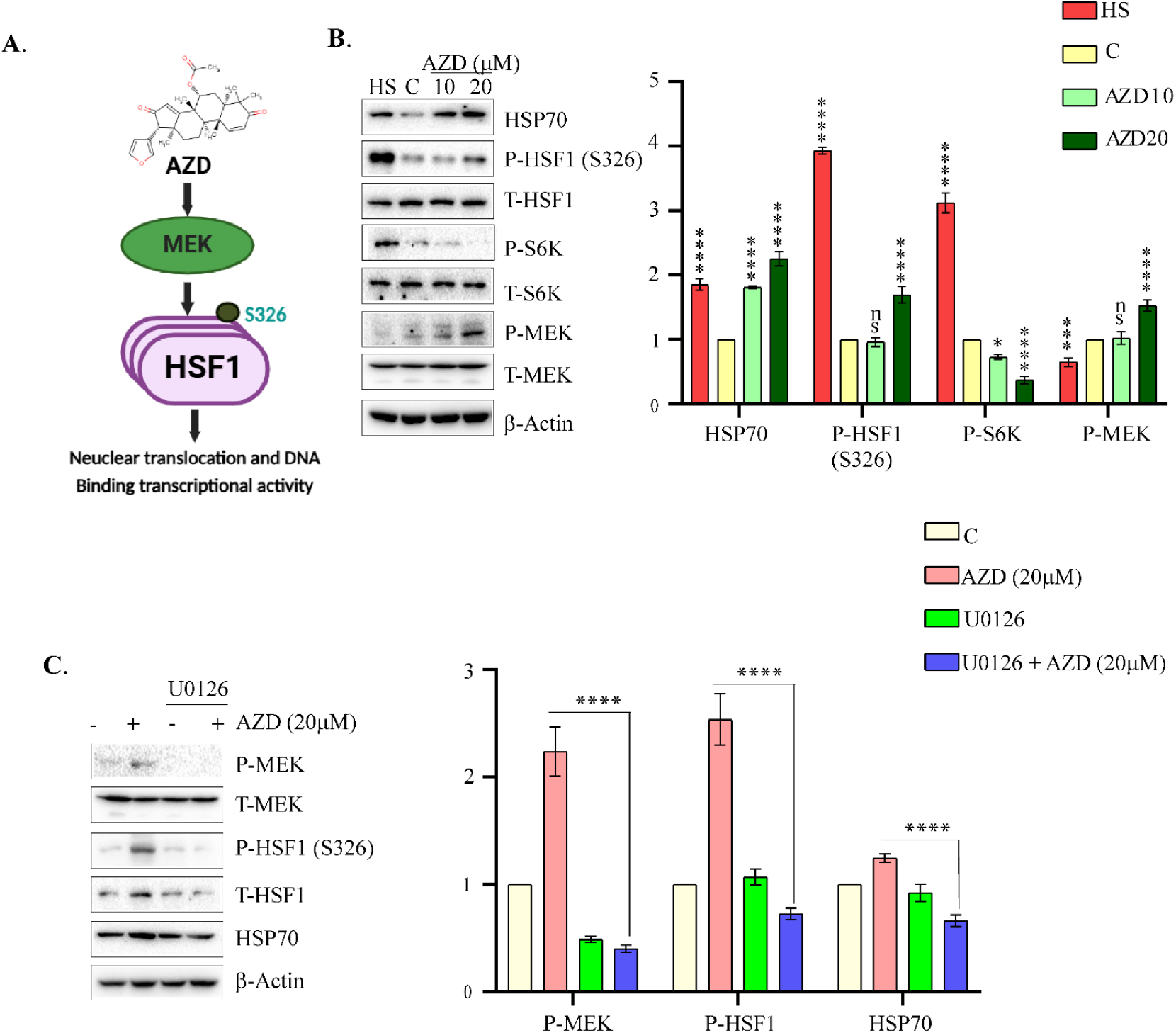
Azadiradione (AZD) induces HSF1 function through MEK activation: *A*, cartoon representing the action of AZD on MEK 1. Only S326 site on HSF1 was shown. *B*, Immunoblots representing the status of indicated proteins in HEK293 cells treated with indicated concentrations of AZD or vehicle (DMSO) for 24 h. β-actin was used as a loading control. Right panel (bar graph) indicates the densitometric analysis of immunoblots using ImageJ software. The levels of the proteins were normalized to loading control β-actin. *C*, Immunoblots representing the effect of MEK inhibitor U0126 on HSF1 activatory phosphorylation. The right panel represents the quantitation of the immunoblots using ImageJ software where the levels of indicated factors were normalized to loading control β-actin. Data represented as mean ± SD of three independent experiments. P value with *P < 0.01 ; ** P< 0.001 and **** P<0.0001 respectively.

Involvement of MEK in AZD function was confirmed by treating the cells with the MEK inhibitor U0126 which abolished S326 phosphorylation and HSF1 activation, resulting in decrease in cellular HSP70 levels [Fig.1*C*][49].

### Azadiradione facilitates clearance of cellular protein aggregates through upregulating proteasome and autophagy pathways

A hallmark of disrupted cellular protein homeostasis associated with aging is the accumulation of protein aggregates, such as those containing polyglutamine (PolyQ) stretches, as seen in Huntington’s disease [50]. Our studies previously revealed that PolyQ-protein aggregation was reduced in AZD-treated cells [36]. Here, we demonstrated that AZD mediates this through activating HSF1, as downregulation of HSF1 by siRNA abrogated this effect in Neuro-2A cells [Figure 2A.i]. The AZD-induced GFP-ataxin 130Q aggregates were reduced in cells both in size and numbers in HSF1 dependent manner (Figure 2A.iii). To better understand we tested if proteasome was involved in the process. We found that AZD significantly increased proteasome activity by dose-dependent upregulation of chymotrypsin activity in the lysates of cells pre-treated with AZD but not with vehicle. In this experiment bortezomib, a proteasome inhibitor was used as a negative control, which, as expected, inhibited the chymotrypsin activity [Figure 2B]. The proteasome activation property of AZD was further tested by measuring the poly-ubiquitinated protein levels in the lysates of cells pre-treated with either the vehicle or AZD, with or without bortezomib. AZD was applied in increasing doses to the bortezomib pre-treated cells, to reveal that AZD treatment could significantly reduce the load of poly-ubiquitinated protein accumulation induced by bortezomib-pre-treatment [Figure 2C].

**Figure 2.**
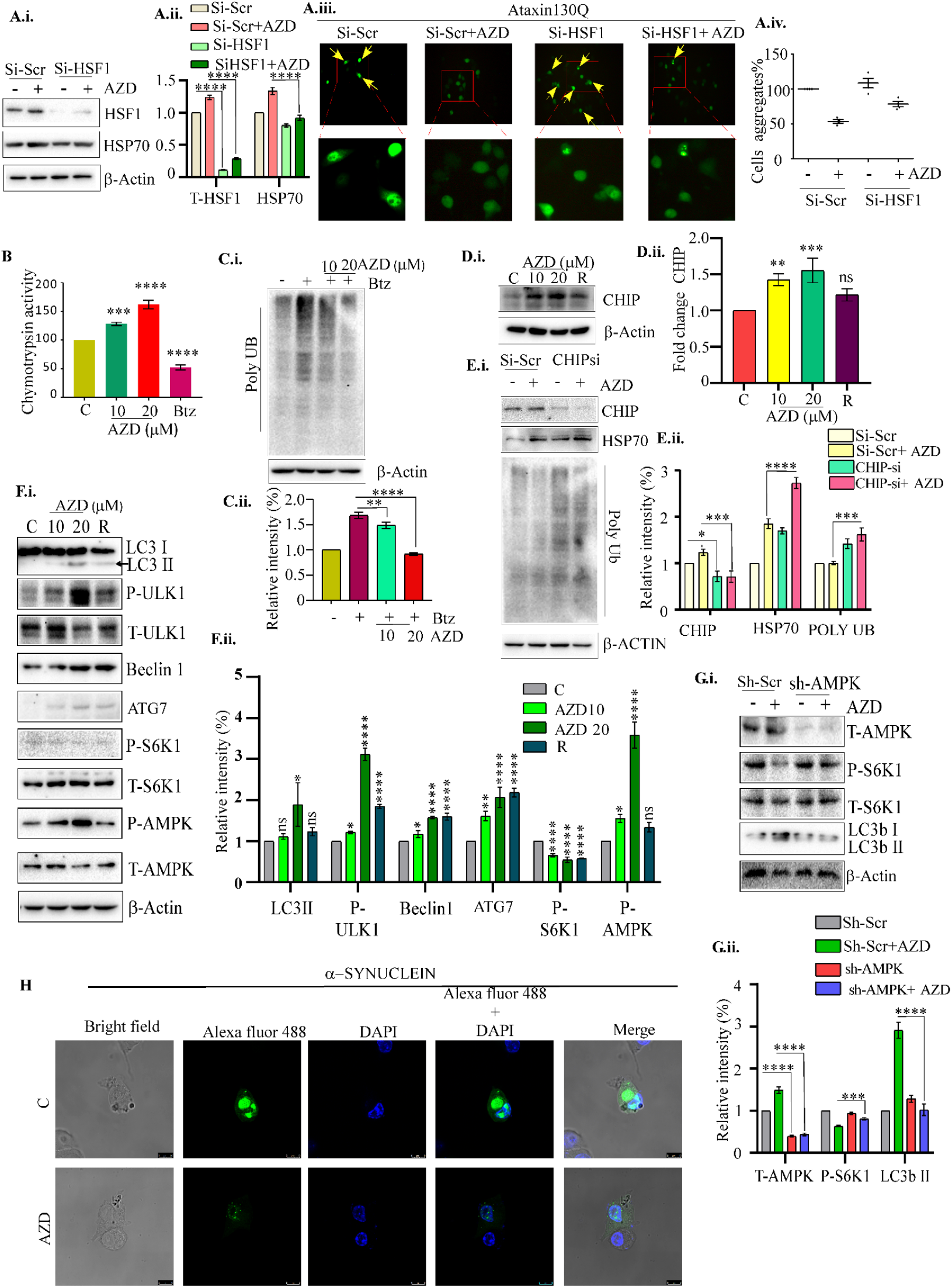
Azadiradione (AZD) upregulates protein quality control machinery in Neuro-2A cells: Left panel (*A*.i.) immunoblots representing the efficacy of siRNA mediated knockdown of HSF1 and HSP70 in cells pre-treated with scrambled siRNA (Si-Scr) or HSF1 siRNA (sh-HSF1). Bar graph indicates the densitometric analysis of immunoblots using ImageJ software (A.II.). The levels of the factors were normalized to levels of loading control β-actin. Data are presented as mean ± SD of three independent experiments. *A*.iii. fluorescence microscopy images representing the HSF1-dependent effect of AZD on cellular aggregation of over-expressed GFP-ataxin130Q protein. The right panel (A.iv.) shows the quantitation of the image using ImageJ software. The aggregates (arrow heads)) counted from three different fields in treated cells in triplicates were plotted. (Data are presented as mean ± SD of three independent experiments.) *B*, bar graph representing 20S proteasome (chymotrypsin) activity in lysates of cells pre-treated with AZD for 24 h. Bortezomib (Btz) was used as a positive control as Proteasome inhibitor. *C*.i., representative immunoblots showing the level of poly-ubiquitinated proteins in extracts of cells pre-treated with AZD. Cells were pre-treated with Bortezomib (Btz 100 nM) for 2 h. β-actin level was measured as an internal loading control. *C*.ii. panel represents the densitometric quantitation of the immunoblots using ImageJ software. The levels of the factors were normalized to levels of loading control β-actin. *D*.i. immunoblots representing the levels of cellular CHIP upon the treatment with AZD. The adjacent right panel (*D*.ii.) represents the densitometric quantitation of the immunoblots using ImageJ software. The levels of these factors were normalized to levels of loading control β-actin. E.i. immunoblots representing the involvement of CHIP on AZD function in cells pre-treated with scrambled siRNA (Si-Shr) or siRNA against CHIP. *E*.ii. panel represents the densitometric quantitation of the immunoblots using ImageJ software. The levels of the factors were normalized to levels of loading control β-actin. *F*.i., immunoblots representing the status of indicated factors in cells pre-treated with AZD. Rapamycin (R, 200 nM) was used as a positive control as an mTORC1 inhibitor. *F*.ii. band intensities in the blot were estimated by densitometric scanning and plotted to compare their expression levels. ß-actin levels were used as internal loading control. *G*.i. immunoblots representing the role of mTORC1 and AMPK in AZD-induced autophagy in cells pre-treated with scrambled shRNA (Sh-Scr) or sh RNA against AMPK. *G*.ii. panel represents the densitometric quantitation of the immunoblots using ImageJ software. The levels of the factors were normalized to the levels of loading control β-actin. *H*. representative fluorescence microscopy images representing the effect of AZD in cells over-expressing α-synuclein double mutants (A30T and A50T, FLAG-epitope tagged). All Data are presented as mean ± SD of three independent experiments. P value with *P < 0.01 ; ** P< 0.001 and **** P<0.0001 respectively.

We also tested the involvement of CHIP (carboxy terminus of Hsc70 interacting protein with E3 ubiquitin ligase property), which links the cellular proteasome system with chaperone proteins through marking misfolded proteins for degradation[31], and noted increased CHIP levels in AZD-treated cells [Fig. 2*D*] as downregulation of CHIP by siRNA abrogated the proteasome activity in these cells [Fig. 2*E*].

Autophagy is another cytoprotective mechanism that facilitates the degradation and recycling of misfolded protein aggregates. Under chronic stress, when the cellular damage or load of misfolded protein aggregates exceeds the repairing capacity of chaperone proteins, autophagy becomes the primary response to promote protein homeostasis[51]. Increased cleavage of LC3-II protein in AZD-treated cells suggested its potential as an autophagy inducer. We also observed that AZD could activate ULK1 function by phosphorylating it at S555, and induce Beclin1 and ATG7, the crucial components of the autophagy process [Fig. 2*F*]. Inhibition of mTORC1 has encouraged us to test the involvement of AMPK/mTOR pathway in AZD treated cells [Fig. 2*F*]. Our results indicate that AZD induces autophagy by modulating the AMPK/mTOR pathway. Because AZD treatment inhibits S6K1 phosphorylation, a downstream target of mTORC1 through enhancing AMPK function by dose-dependent increase of AMPK phosphorylation at T172 [Fig. 2*F*]. We noted that shRNA-mediated downregulation of AMPK abrogated AZD activity on mTORC1 and LC3-II cleavage Role of AZD in improving the cellular protein quality was further tested in another protein aggregation model apart from PolyQ. We observed that AZD did reduce aggregates of alpha-synuclein double mutant (A30P/A50T), overexpressed in Neuro-2A cells (Fig. 2*H*). Aggregation of alpha-synuclein mutants in the brain is a signature of the development of Parkinson’s disease (PD)[41, 52].

### Azadiradione controls AKT-FOXO and antioxidant pathway

FOXO transcription factors integrate a wide range of signaling pathways in response to various cellular conditions[53]. Nutritional stress, such as calorie restriction, can upregulate these transcription factors by modulating them through diverse post-translational modifications[54]. The activation of AMPK and inhibition of mTORC1 suggest that AZD influences the cellular calorie restriction pathway (Fig. 2*F*). To explore this, we examined the status of FOXO following AZD treatment and found that AZD suppresses the phosphorylation of FOXO3 at S253. AKT-mediated phosphorylation of FOXO3 at S253 typically leads to its nuclear exclusion[55]. However, AZD-mediated inhibition of AKT promotes FOXO3 dephosphorylation, enabling its nuclear translocation and subsequent activation of target genes, including SOD and catalase (Fig. 3*A, B*).

**Figure 3.**
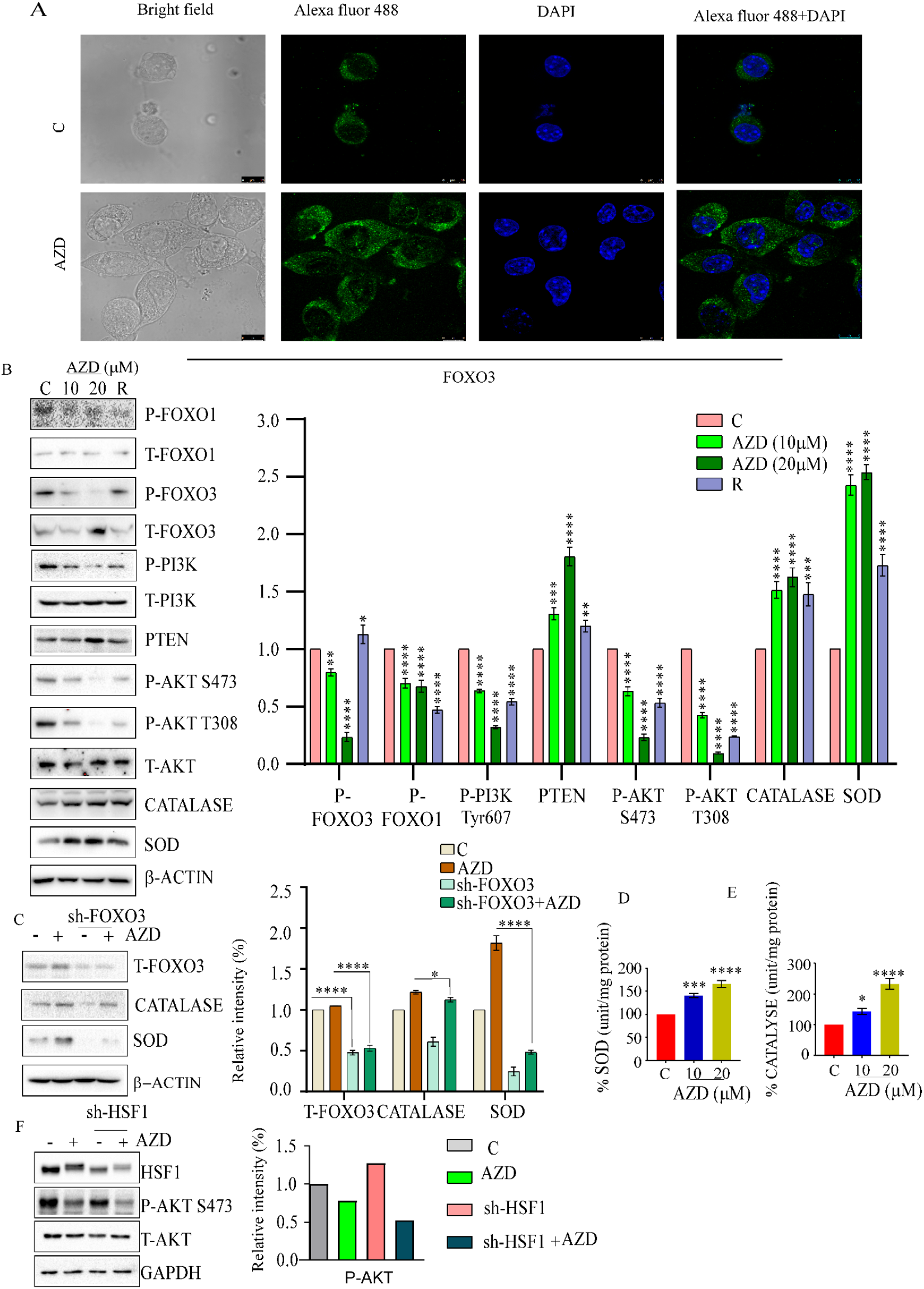
Azadiradione (AZD) modulates Insulin signalling via AKT-FOXO pathway: *A*, Immunofluorescence images representing the translocation of FOXO3 to the nucleus upon AZD treatment. Cells were treated with 20 µM of AZD for 18 h. *B*, immunoblots representing the status of FOXO1, FOXO3a and their targets SOD and Catalase, and AKT in lysates of Neuro-2A cell pre-treated with AZD for 24 h. The right panel represents the densitometric quantitation of the immunoblots using ImageJ software. The levels of the factors were normalized to levels of loading control β-actin. *C*, immunoblots representing the dependence of AZD function on FOXO3a for SOD and Catalase as downregulation of FOXO3 by shRNA but not by scrambled shRNA (Sh-Scr). *D*-*E*, bar graphs representing the SOD and Catalase activity measured in lysates of cells pre-treated with AZD for 24 h. The levels of the enzymes were normalized to their levels in mock treated cells. *F*, immunoblots representing the dependence of AZD functions on AKT phosphorylation in the presence of Sh-HSF1 and scramble HSF1 (Sh-Scr). The right panel represents the densitometric quantitation of the immunoblots using ImageJ software. The levels of the factors were normalized to levels of loading control β-actin. All Data are presented as mean ± SD of three independent experiments. P value with *P < 0.01 ; ** P< 0.001 and **** P<0.0001 respectively.

Additionally, in the same cells, similar to FOXO3, the phosphorylation levels of FOXO1 (S256) and AKT (S473, T308) were also downregulated (Fig. 3*B*)[56, 57]. We further confirmed that the induction of SOD and catalase protein levels by AZD was compromised in cells pre-treated with FOXO3 siRNA (Fig. 3*C*). Moreover, AZD not only increased the protein levels of SOD and catalase but also enhanced their enzymatic activity in lysates of cells pre-treated with AZD (Fig. 3 *D, E*). These effects of AZD are independent of HSF1, as HSF1 downregulation did not alter AZD-induced changes (Fig.3*F*). Recent reports also suggest that AZD mimics the function of SOD[58]. Finally, the inhibition of PI3K, upstream of AKT, indicates that AZD targets the PI3K-AKT signaling pathway upstream of AKT, potentially through P-TEN-mediated mechanisms[59].

### Azadiradione ameliorates Parkinson’s disease in mice

We tested the effect of AZD on a mouse model of Parkinson’s disease (PD) created using MPTP (1-methyl-4-phenyl-1,2,3,6-tetrahydropyridine) [41]. In astrocytes, MPTP is converted into a toxic form, MPP+, which kills dopaminergic neurons to result in developing PD-like symptoms[60]. In PD, neurons die due to alpha-synuclein clumping into toxic aggregates in the substantia nigra. Heat shock proteins are known to help stabilize alpha-synuclein, reducing its aggregation and toxicity[61] .Our earlier studies showed that increasing HSF1 level by AZD can help improve Huntington’s disease in mice[38].

AZD was given every other day to Swiss Albino mice for 20 days to two groups of Swiss Albino mice (n=5), with one group pre-treated with MPTP [Figure 4A]. We also included control groups (n=5 each) pre-treated with either DMSO or AZD alone. During the experiment, we tracked changes in behavior (grip strength and steps patterns) and body weight. AZD-treated mice regained their lost gripping ability and body weight over time, while MPTP-treated mice did not [Fig. 4B, C]. AZD-treated mice showed significant improvement in their step counts compared to MPTP-only mice [Fig. 4D, E]. Notably, AZD did not cause any visible negative effects on healthy mice [Fig. 4B, C].

**Figure 4.**
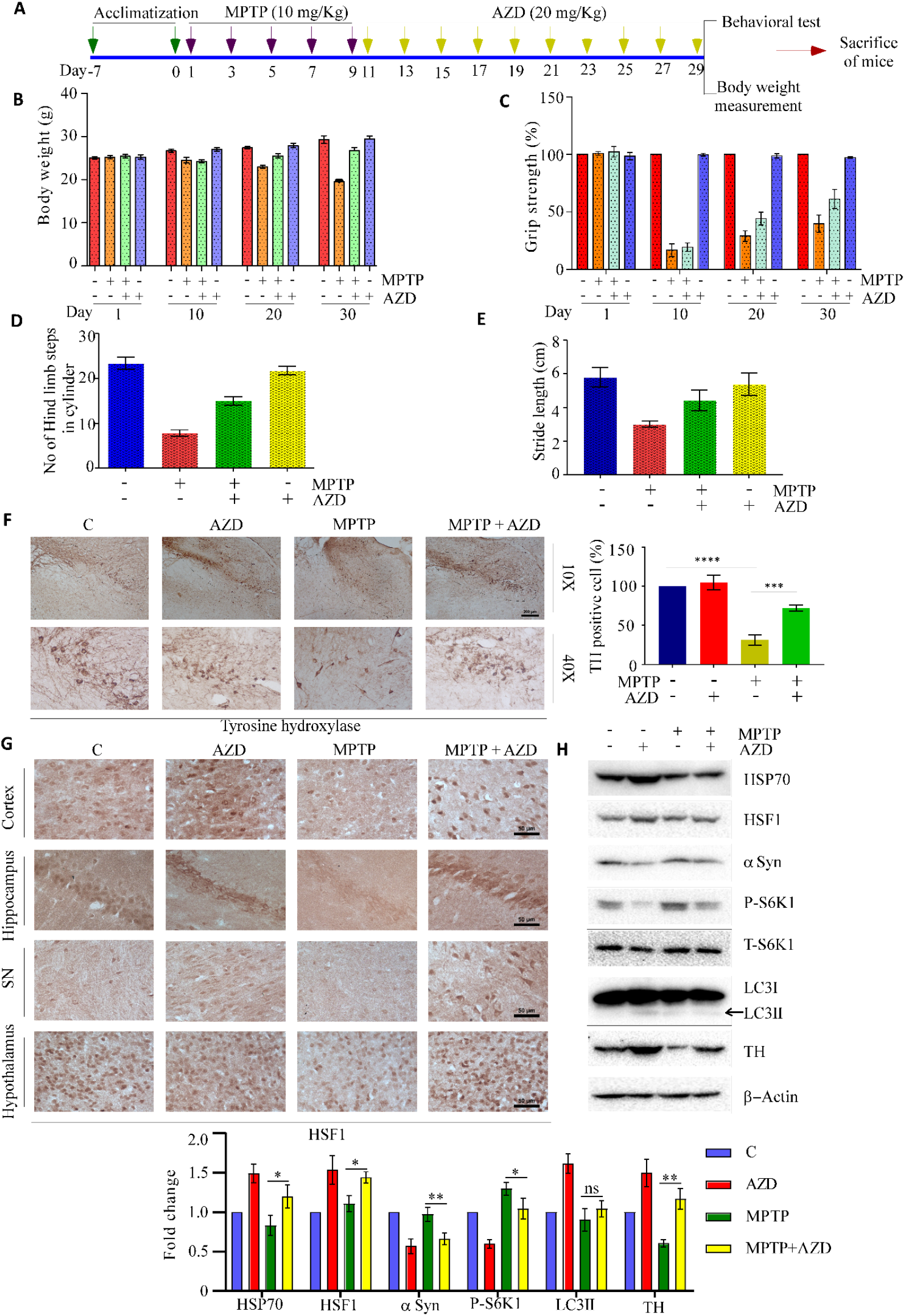
Azadiradione (AZD) ameliorates Parkinson’s disease in MPTP-induced mice model: *A*, schematic overview of experimental design. MPTP and Azadiradione (AZD) concentrations used are indicated. *B*-*C*, bar diagrams representing the recovery of body weight and grip strength in MPTP-treated mice upon AZD treatments. *D-E*, bar diagrams representing the recovery of hind limb step in cylinder and stride length measured in MPTP-treated mice upon AZD treatments. *F*, immunostained images representing tyrosine hydroxylase (TH) levels in substantia nigra pars compacta of mice treated as indicated. The right panel represents the quantitation of the staining as a bar graph on the right. Bar, mean ± SD (n = 5), **** P < 0.0001. *G*, immunostained images representing HSF1 levels in different parts of brain of mice treated as indicated. *H*, immunoblots representing the levels of indicated proteins in lysates isolated from the brains of mice treated as indicated. The right panel represents the densitometric quantitation of the immunoblots using ImageJ software. The levels of the factors were normalized to levels of loading control β-actin. All Data are presented as mean ± SD of three independent experiments. P value with *P < 0.01; **P < 0.001.

AZD helped restore tyrosine hydroxylase (TH) levels, a marker of dopaminergic neurons, in the substantia nigra compared to the MPTP-only group [Fig. 4F]. Immunohistochemistry (IHC) showed that AZD increased HSF1 levels in different brain regions, including the cortex, hippocampus, substantia nigra, and hypothalamus [Fig. 4G]. Finally, immunoblot analysis of brain lysates confirmed that AZD increased HSP70, TH, and LC3-II protein levels while reducing S6K1 and alpha-synuclein levels [Fig. 4H].

### Azadiradione extends healthy lifespan in fruit fly

Upregulation of protein quality control mechanism correlated well with extension of lifespan [62, 63]. Here, we tested the effects of AZD on lifespan extension and age-related physiological changes in the Drosophila model at non-toxic AZD doses. We supplemented growth media with AZD concentration that did not affect the eclosion rate or the egg lying capacity of the flies [Fig. 5B, C].

**Figure 5.**
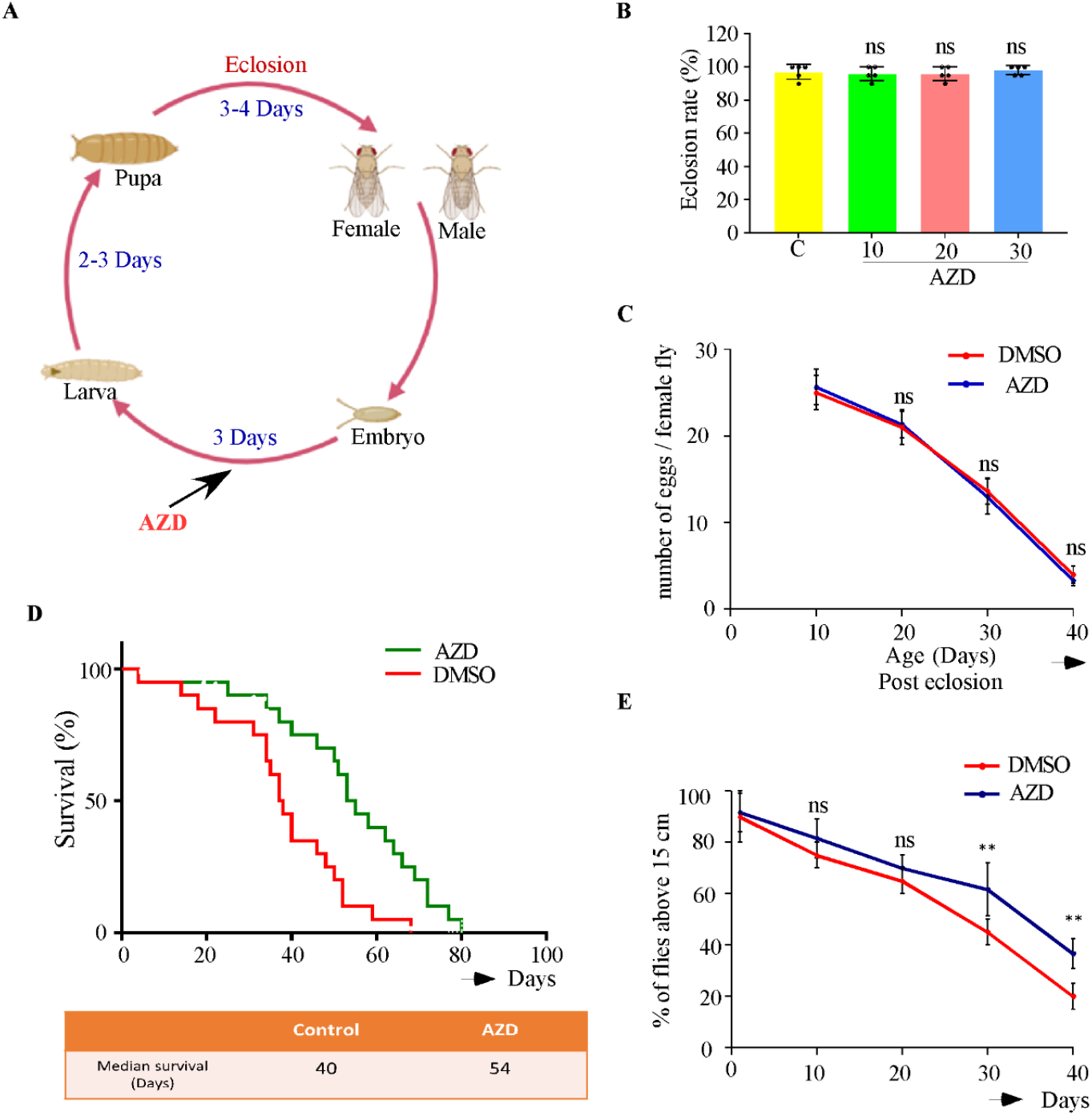
Azadiradione (AZD) extends life span in Fruit fly model without compromising their fertility and physical activities: *A*, schematic diagram showing the *Drossophila melanogaster* life cycle and AZD treatment protocol. Fruit flies were maintained at 25ºC with 12 hours light/dark cycle and grown on media during 1^st^ instar larvae stage. *B*, bar diagram representing well-tolerance of the fruit flies of growth media as measured their eclosion rate, supplemented with AZD as indicated. *C*, line graphs representing reproductive fitness (as egg laying ability) status of age-matched flies maintained on growth media supplemented with AZD or DMSO. *D*, Kaplan-Meier survival curve of flies fed on media supplemented with AZD or DMSO. Fly viability was scored over their full lifetime, using 20 fly per vial. *E*, negative geotaxis assay of the same samples obtain at 10 days, 20 days, 30 days and 40 days. All Data are presented as mean ± SD of three independent experiments.

Our results suggested that AZD-fed flies acquired extension of their lifespan compared to their counterparts maintained on growth media supplemented with the vehicle (DMSO) only (control flies). As revealed the median survival of AZD fed flies was about 54 days compared to that of 40 days of the control flies [Fig. 5D]. Our studies shows that AZD increases around 46% lifespan increases compared to control DMSO fed flies. (More data Set of life span experiment presented in supplementary figure 1) The vehicle-treated group showed a decline of their flying capacity over time. In striking contrast, AZD feeding could suppress the age-associated decline in their flying capacity compared to control flies as these flies did not lose their flying as well as their egg laying capacity [Figure 5E].

### Azadiradione and heat shock alter the expression of distinct and overlapping sets of cellular genes revealed by transcriptome analysis

We conducted a transcriptome analysis via RNA-seq in Neuro 2A cells to comprehensively examine the global gene expression changes upon AZD as well as HS treatment. We have included differentially expressed genes with p-value threshold < 0.05 for analysis. Only the normalized values after adjustment with mock treated cells were included in calculation. Some of the genes in these categories are listed in (Supplementary table 1, Table 2). Gene Ontology (GO) analysis of differentially expressed genes uncovered multiple gene sets associated with HSR, autophagy, proteasome or insulin signalling pathway [Fig. 6*D*]. Many genes associated with autophagy, proteasome, HS and insulin signaling pathway responded differently in AZD and HS treated cells (Fig. 6*D-G*). Comparative expression analyses of more than 5000 genes revealed that, while 1767 genes were influenced by both AZD and HS, 984 and 2602 genes were uniquely influenced by AZD and HS, respectively (Fig. 6*C*). Thus, our transcriptomic analysis demonstrated that AZD controls expression of genes associated with various stress response pathways, as anticipated.

**Figure 6.**
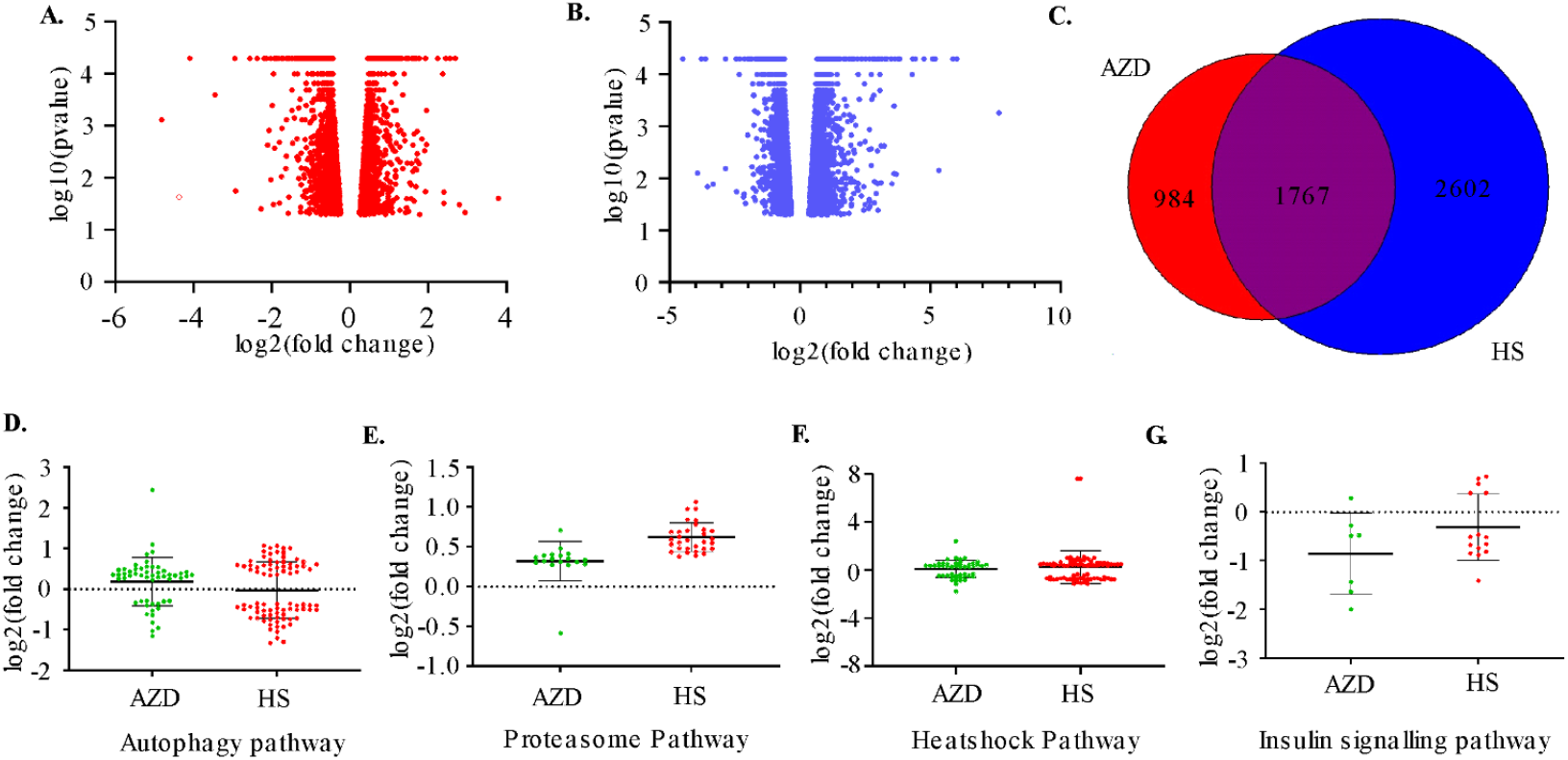
Transcriptomic analysis of cellular genes upon heat shock and AZD treatment: (*A, B,*) volcano plot representing expression of genes modulated upon the treatment with AZD or heat shock (HS) in Neuro-2A cells. *C*, venn diagram representing the group of genes regulated by either AZD, HS or both AZD and HS. Sets of differentially expressed genes under p-value threshold < 0.001 were selected for analysis is reported. *D*-*G*, representative graphs showing the change in the expression of genes of the indicated pathways in AZD or HS treated Neuro-2A cells.

## Discussion

We reported earlier that AZD mitigates cellular toxicity caused by protein aggregation by activating of HSF1 [36, 38] .Our previous observation of AZD-induced DNA binding by HSF1 prompted us to analyze the phosphorylation status of HSF1 at the S326 residue, a known hallmark of HSF1 activation [26, 48]. Various kinases, including mTORC1, AKT, p38, DYRK2, and MEK1 were implicated in the phosphorylation at this site [28, 48, 64]. Tang et al. demonstrated that HS-induced RAS/MAPK signaling facilitated MEK-mediated S326 phosphorylation through direct interactions with HSF1 [28]. We found that AZD promotes this phosphorylation of HSF1 via MEK activation, while inhibiting mTORC1 (Fig. 1*B*) although further studies will be required to test the involvement of other kinases in this process. These observations suggest that MEK activation by AZD not only enhances DNA-binding affinity of HSF1 but also stabilizes HSF1 by preventing its proteasomal degradation. Notably, AKT activation (phosphorylation) is also inhibited in AZD-treated cells (Fig. 3*B*,*F*).

Earlier we demonstrated that AZD ameliorates protein aggregation-induced vision loss in fruit flies [36]and Huntington’s disease in mice [38]. AZD activates cellular E3 ubiquitin ligase, CHIP to facilitate the degradation of misfolded proteins by the proteasome (Fig. 2*D*)[38]. Furthermore, AZD could ameliorate PD symptoms and pathology in mice by activating both the proteasome and autophagy pathways (Fig. 2). These results are in agreement with our whole-genome transcriptome analysis which suggested that like HSR, AZD upregulates genes associated with both the proteasome and autophagy pathways (Supplementary Table 1, TABLE 2). Altogether these results highlighted the therapeutic potential of AZD for NDs through simultaneously modulating multiple PQC pathways, albeit future tests will necessary to help further understanding of this hypothesis.

Free radical-induced oxidative stress is considered as a major cause of damage to cellular macromolecules leading to aging [65, 66]. FOXO family transcription factors such as FOXO3 and FOXO1 are implicated in maintaining protein homeostasis under oxidative stress [67]. FOXO3 activity is controlled by modifications like phosphorylation through regulating its subcellular location [57, 68, 69]. AKT phosphorylates FOXO3 at multiple sites, moving it from the nucleus to the cytoplasm and blocking its function as a transcription factor[68, 70]. This study revealed that AZD can modulate the PI3K/AKT/FOXO3 axis to upregulate cellular antioxidant pathways via inducing SOD and Catalase activities in consistent with its observed lifespan extension function (Figs. 3 & 5). FOXO can also influence cellular autophagy [71]. Preliminary effect of AZD on insulin signalling pathway supports lifespan extension as well (Fig. 6*G*). Earlier, AZD was reported to activate HSF1 functions with little interruption on cellular oxidative status, HSP90 or proteasome [36]. HSF1-independent inhibition of AKT suggested modulation of additional cellular pathways by AZD which might not be surprising as natural products were shown to modulate multiple cellular pathways (Fig. 3*F*)[72]. Nevertheless, abrogation of AKT function favours the idea that HSF1 activator AZD does not support proliferative cellular pathway correlated with cancer. AZD-loaded liposome was shown to inhibit triple negative breast cancer in mice[73]. In fact, in increasing body of evidence suggested an inverse correlation between many types of cancer and neurodegenerative diseases such as PD and Alzheimer’s disease (AD)[74]. That is, PD and AD patients are relatively less likely to have certain types of cancers [75-77].

AZD-mediated autophagy induction involving cellular AMPK activation and TORC1 inhibition correlated well with amelioration of protein aggregation induced toxicity and life span extension (Figs. 2*F* and 5). AMPK can activate autophagy directly by phosphorylating ULK/ATG1 complex or through inhibiting mTORC1 through phosphorylating Raptor or TSC2 in the mTORC1 complex [78].

Unlike many other small-molecule HSF1 activators, AZD possesses unique properties—it upregulates cellular PQC pathways without affecting the healthy mice. Notably, AZD antagonises function of AKT, a kinase that promotes cancer cell growth, through an HSF1-independent mechanism, suggesting there is much more to understand about its cellular function. Altogether, these findings highlight the importance of continued research into AZD’s cellular functions—not only for its therapeutic potential in PD but also for its possible relevance in other NDs and for gaining deeper insights into cellular signalling pathways.

## Author contributions

N.D., H. R. S., S. G., U. C. V. K. N. data curation; N.D., H. R. S formal analysis; N.D., H. R. S. and S. G., U. C., V. K. N., A. K. M. and M. P. validation; N. D., H. R. S, S. M., and M. P. investigation; N.D., H. R. S., and M. P. visualization; N.D., H. R. S., and S. G., U. C., V. K. N. methodology; N.D., and H. R. S., writing-original draft; A. K. M. and M. P. Resources; S. M., and M. P. conceptualization; M. P. supervision; M. P. project administration; A. K. M., and M. P. writing-review and editing.

## Acknowledgments

We thank DBT, SERB and Bose Institute for funding, and faculty members of Division of Molecular Medicine, now named Department of Biological Sciences of Bose Institute for their continuous help to carry out this work. We gratefully acknowledge the help provided by Dr. Kuladip Jana during the animal experiment.

