## Supplementary material for "HSF1 Activator Azadiradione Ameliorates Parkinson’s Disease and Extends Lifespan in Preclinical Models: Analysis of Underlying Molecular Mechanism": SUPLIMENTARY DATA

Supplementary figure

A.

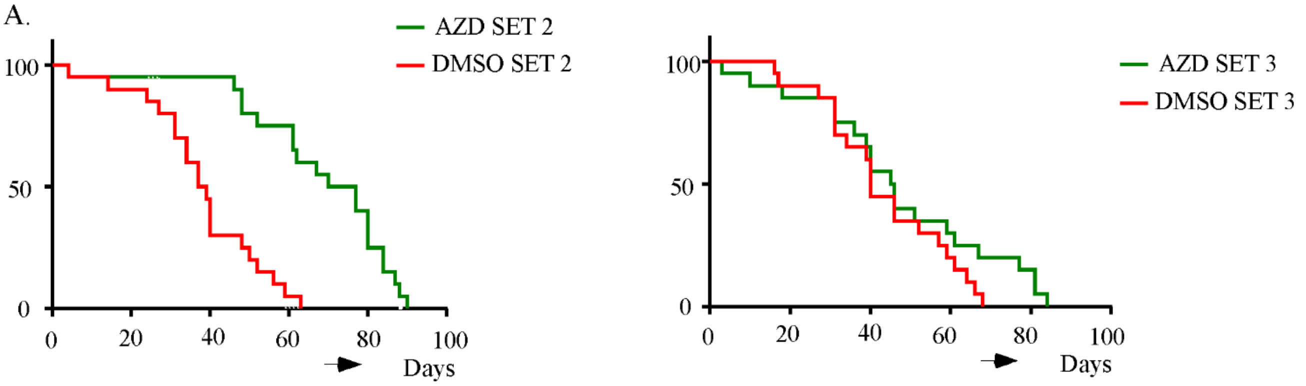

| No Of SET | Median survival of AZD fed flies | Median survival of DMSO fed flies |
|---|---|---|
| SET 1 | 54 | 40 |
| SET 2 | 73.5 | 38 |
| SET 3 | 45.5 | 40 |

Supplementary table 1

**TABLE 1**

**List of Antibodies**

| **Antibody** | **Source** | **Catalogue No** |
|---|---|---|
| HSF1 | BioBharati # | BB-AB0220 |
| Phospho- HSF1 | Abcam | ab76076 |
| Actin | Abcam | ab49900 |
| HSP70 | BioBharati | # BB-AB0210 |
| Phospho-MEK1/2 (Ser217/221) Antibody | Cell signaling technology | #9121 |
| MEK1/2 Antibody | Cell signaling technology | #9122 |
| AMPK | Cell signaling technology | #2532 |
| Phospho- AMPK | Cell signaling technology | #2535 |
| S6K | Cell signaling technology | #9202 |
| Phospho-S6K | Cell signaling technology | #9205 |
| LC3b | Cell signaling technology | #3868 |
| AKT | Cell signaling technology | #9272 |
| Phospho-AKT | Cell signaling technology | #9271 |
| ATG7 | Cell signaling technology | #8558 |
| Phospho-ULK1 (Ser555) (D1H4) Rabbit mAb | Cell signaling technology | #5869 |
| ULK1 (D8H5) Rabbit mAb | Cell signaling technology | #8054 |

| | | |
|---|---|---|
| Beclin-1 Antibody | Cell signaling technology | #3738 |
| Ubiquitin | Cell signaling technology | #3933 |
| PTEN (138G6) Rabbit mAb | Cell signaling technology | #9559 |
| CHIP | Santa Cruz Biotechnology | #SC133083 |
| Tyrosine hydroxylase | Cell signaling technology | #2792 |
| Phospho-FoxO1 (Ser256) Antibody | Cell signaling technology | #9461 |
| FoxO1 (C29H4) Rabbit mAb | Cell signaling technology | #2880 |
| Phospho-FoxO3a (Ser253) (D18H8) Rabbit mAb | Cell signaling technology | #13129 |
| FoxO3a (75D8) Rabbit mAb | Cell signaling technology | #2497 |
| Phospho-PI3 Kinase p85 (Tyr458)/p55 (Tyr199) (E3U1H) Rabbit mAb | Cell signaling technology | #17366 |
| PI3 Kinase p110α (C73F8) Rabbit mAb | Cell signaling technology | #4249 |
| Goat anti-rabbit IgG-HRP | Santa Cruz Biotechnology | sc2004 |
| Goat anti-mouse IgG-HRP | Santa Cruz Biotechnology | sc2005 |
| α-Synuclein | Cell signaling technology | #4179 |
| Goat anti-mouse IgG-Alexa Fluor 488 | Thermo Scientific | #A32723 |
| Goat anti-rabbit IgG-Alexa Fluor 488 | Thermo Scientific # | A-11008 |

**TABLE 2**

IMPORTANT GENES INDUCED BY AZD

| AUTOPHAGY | HEATSHOCK RESPONSE | PROTEASOME | INSULIN SIGNALLING PATHWAY |
|---|---|---|---|
| Hmox1 | Hmox1 | Psmd4 | Irs1 |
| Sqstm1 | Hsp90ab1 | Psmb3 | Pik3r1 |
| Atg101 | Hspa5 | Psma5 | Igfbp5 |
| Atg12 | Hspa8 | Psma1 | Igf1r |
| Pink1 | Hsp90aa1 | Dnajb2 | Sos1 |
| Stub1 | Hsp90ab1 | Psma7 | Grb10 |
| Bag3 | Stub1 | Psmc2 | Map2k1 |
| | Dnaja1 | Psmd7 | |
| | Sod1 | Psmd3 | |
